# Gene buddies: Linked balanced polymorphisms reinforce each other even in the absence of epistasis

**DOI:** 10.1101/220897

**Authors:** Jacob A Tennessen

## Abstract

The fates of genetic polymorphisms maintained by balancing selection depend on evolutionary dynamics at linked sites. While coevolution across linked, epigenetically-interacting loci has been extensively explored, such supergenes may be relatively rare. However, genes harboring adaptive variation can occur in close physical proximity while generating independent effects on fitness. Here, I present a model in which two linked loci without epistasis are both under balancing selection for unrelated reasons. Using forward-time simulations, I show that recombination rate strongly influences the retention of adaptive polymorphism, especially for intermediate selection coefficients. A locus is more likely to retain adaptive variation if it is closely linked to another locus under balancing selection, even if the two loci have no interaction. Thus, two linked polymorphisms can both be retained indefinitely even when they would both be lost to drift if unlinked. Such clusters of mutually reinforcing genes may underlie phenotypic variation in natural populations. Future studies that measure selection coefficients and recombination rates among closely linked genes will be fruitful for characterizing the extent of this phenomenon.

## Introduction

Balancing selection is an evolutionary process that maintains more than one allele at a locus for longer than would be expected under genetic drift alone (Fijarczyk and Babik 2015). Numerous examples of balanced polymorphisms are well known, related to infectious disease (Pasvol et al. 1978; Hughes and Nei 1988; Tennessen and Blouin 2008), reproduction (Wright 1939; Käfer et al. 2017), and other traits (reviewed in Llaurens et al. 2017). Balancing selection can operate via several distinct mechanisms, including overdominance or heterozygote advantage, frequency-dependent selection, or spatiotemporally varying selection (Tennessen and Blouin 2008). While early population geneticists proposed widespread balancing selection to explain polymorphisms (Lewontin and Hubby 1966), this view fell out of favor with the rise of the neutral theory (Kimura 1983). However, without validating the pan-adaptationism of earlier decades, the importance of balancing selection is being increasingly recognized (Sellis et al. 2011; Key et al. 2014; Delph and Kelly 2014).

For an adaptive polymorphism to survive, balancing selection must overcome genetic drift (Sutton et al. 2011), which is primarily determined by the effective population size (*N_e_*). When *s*, the selective advantage of a genotype, is much less than 1/*N_e_*, the polymorphism is effectively neutral and it is eventually lost, whereas if *s* is much greater than 1/*N_e_*, the effects of selection will govern evolution (Ohta 1992). Thus, when *s* is close in magnitude to 1/*N_e_*, the opposing forces of balancing selection and drift nearly cancel out, and the fate of a balanced polymorphism may be more heavily influenced by other factors, including selection acting on linked sites. Linkage disequilibrium (LD) between physically adjacent loci has been recognized as an important factor determining evolutionary outcomes, and thus a haplotype block of several loci in LD, rather than an individual locus, may be thought of as the unit of selection (Casillas and Barbadilla 2017). The extent of LD between two linked loci depends on the recombination rate (*r*), *N_e_*, and other evolutionary forces. For balancing selection, this haplotype framework raises the question of if, and how, genetic variation at one locus affects the probability that adaptive variation will be maintained at linked loci.

One obvious mechanism is epistasis: if only certain combinations of alleles at two linked loci yield high fitness, then allele frequencies at one locus will depend on those at the other locus, resulting in epistatically interacting “supergenes” (reviewed in Thompson and Jiggins 2014; Llaurens et al. 2017). Intuitively, if the fitness effect of one locus depends on the genotype at a second, linked locus, then not only might selection favor LD between these loci, but the strength of selection on the resulting multi-locus haplotype may be greater than it would be for either locus in isolation. In deterministic models, linkage only affects equilibrium allele frequencies when there is epistasis (Lewontin and Kojima 1960; Nagylaki 2009). Thus, explanations for clusters of balanced polymorphisms such as sex chromosomes (Bergero and Charlesworth 2009), the major histocompatibility complex (Slatkin et al. 2000; Trowsdale 2002; Makino and McLysaght 2008), or generalized quantitative trait architecture (Karlin and Feldman 1970; Draghi and Whitlock 2014) often implicitly assume epistasis. Epistasis is ubiquitous in genotype/phenotype relationships, and evidence of epistasis between closely-linked loci has been observed in some cases (e.g. Rice 1987; Gregersen et al. 2006). However, in many systems there is little empirical evidence that co-adaptation among linked genes maintains polymorphism (Ohta 1982; Navarro and Barton 2002; Wegner 2008; Corbett-Detig and Hartl 2012). In particular, while sex chromosomes are the most striking examples of supergenes both in their taxonomic ubiquity and their extreme heteromorphism, the degree to which epistasis (i.e. sexual antagonism) shapes sex chromosome evolution remains contentious (Fry 2009; Ironside 2010; Pennell et al. 2015; Branco et al. 2017).

If previous work has overemphasized the importance of epistasis in shaping linked adaptive polymorphisms, it is worth exploring the dynamics of linked polymorphisms with no functional relationship. In finite populations, genetic drift may prevent weakly selected loci from reaching equilibrium conditions, and in these cases, linkage can have a nontrivial effect. Thus, polymorphisms maintained by balancing selection could become clustered in the genome not because they interact, but because selection acts more efficiently on multi-locus haplotypes than on individual loci. Here I explore the extent to which polymorphism at a locus under balancing selection depends on other independent instances of balancing selection, acting for unrelated reasons on linked loci.

## Methods

In order to test whether balanced polymorphisms can show mutual reinforcement in the absence of epistasis, I used forward-time simulations. I simulated the evolutionary dynamics of two linked polymorphisms under balancing selection in a finite population with no epitasis. The fitness of a diploid individual was 1 for double homozygotes, and larger by *s* for each heterozygous locus, such that double heterozygotes had a fitness of 1 + 2*s*. Thus, the model was consistent with overdominance, which is mathematically equivalent to some forms of frequency-dependent selection (Takahata and Nei 1990). Following the format of Lewontin and Kojima (1960), this model conforms to the formula

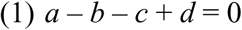

where *a* is the fitness of double homozygotes (here 1), *b* and *c* are the fitnesses of each single heterozygote (here both 1 + *s*) and d is the fitness of double heterozygotes (here 1 + 2*s*). Thus, there is no epistasis as defined by Lewontin and Kojima (1960).

I set the initial frequencies of all four haplotypes to 25%. That is, both polymorphisms occurred at 50% frequency and were not in linkage disequilibrium with each other. I set the population size at 1000, thus *N_e_* = 2*N* = 2000. I set s to various values between 0.002 and 0.01, thus *N_e_s* ranged from 4 to 20. I set *ρ*, the population-scaled recombination rate (=4*N_e_r*), to 0 (perfect linkage), infinite (no linkage), or various values ranging from 0.005 to 20. Simulations were run forward in time for 100,000 (= 50 *N_e_*) generations, with 1000 replicates of each parameter combination. I recorded how often one or both polymorphisms were retained. For all combinations of parameters, I recorded the median time to fixation (MTF) for a polymorphism. In order to test for a significant effect of *ρ* on MTF, I ran linear regression of log(MTF) as a function of log(*ρ*). I also recorded the number of simulations (out of 1000 replicates) for which both polymorphisms were retained for all 50 *N_e_* generations, and I ran linear regression of the log of this value plus one (potential range log(1) to log(1001)) as a function of log(*ρ*). All simulations can be run using a Perl script available at https://github.com/jacobtennessen/GeneBuddies.

## Results

The efficacy of selection varied widely depending on model parameters. Allele frequencies typically fluctuated substantially, eventually fixing (Figure 1), but the fates of balanced polymorphism depended strongly on recombination rates, especially at intermediate selection strengths.

**Figure 1.**
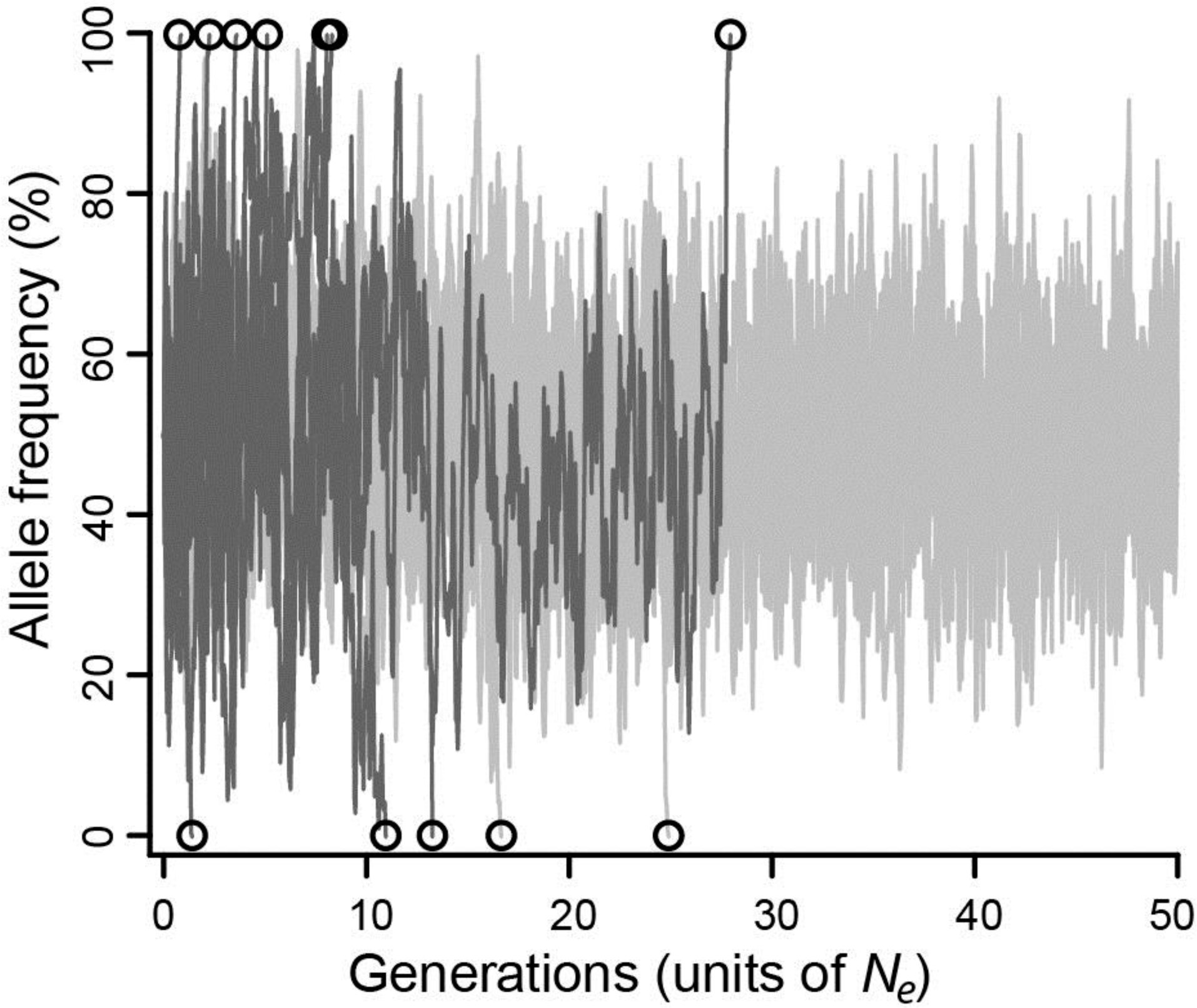
Allele frequencies across 100,000 generations (= 50 *N_e_*) at a locus under balancing selection linked to another locus under balancing selection, with no epistasis. 20 simulations are depicted, all with intermediate selection (*N_e_s* = 10), of which 10 have no recombination (light grey), and 10 have infinite recombination (dark grey). Circles mark the moment of fixation. With perfect linkage (light grey), approximately 80% of simulated loci retain their polymorphism for the duration of the simulation (8 out of 10 do so here), while under negligible linkage (dark grey), nearly all loci fix well before 50 *N_e_* generations (all 10 do so here).

When selection was weak (*N_e_s* ≤ 5), polymorphism was lost relatively quickly in 2 to 7 *N_e_* generations regardless of recombination rate, whereas if selection was strong (*N_e_s* ≥ 15), polymorphism was typically retained for the duration of the simulated 50 *N_e_* generations. However, for all values of *N_e_s*, lower values of *ρ* were associated with greater retention of polymorphism (Figure 2). Specifically, for *N_e_s* values of 12 and under, there was a significantly negative relationship between *ρ* and MTF (*P* < 0.05; Table 1). The strongest effect was observed for *N_e_s* of 8, such that reducing *ρ* by 50% resulted in a 29% increase in MTF (*P* < 10^-5^; Table 1), and thus MTF varied by an order of magnitude depending on *ρ*. A similar trend occurred for higher values of *N_e_s*, but since polymorphism was typically retained for all 50 *N_e_* generations even at high *ρ* values, the effect could not be rigorously quantified.

**Figure 2.**
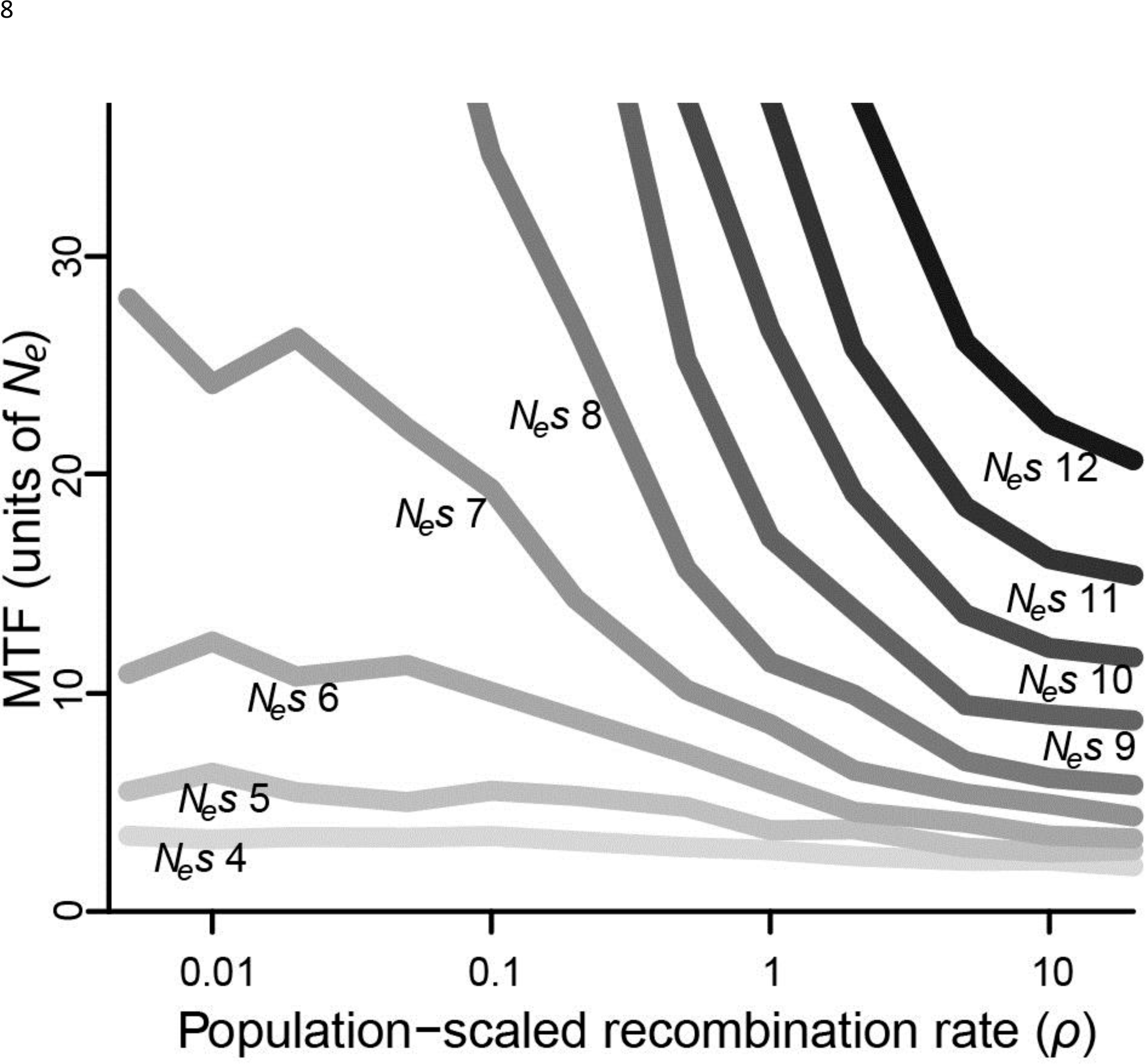
Median time to fixation (MTF) for a polymorphism linked to another polymorphism, for varying values of selection (*N_e_s* ranging from 4 to 12) and recombination (*ρ* ranging from 0.005 to 20). Fixation takes longer when recombination is low, although the magnitude of this effect depends on the strength of selection. Results for *N_e_s* values larger than 12 are not shown, as most polymorphisms endured for all 50 *N_e_* generations across most values of *ρ*.

**Table 1.** Values of *N_e_s* for which *ρ* had a significant effect on median time to fixation (MTF) over 50 *N_e_* generations.

| $N_e s$ | % increase in MTF if $\rho$ is halved | $P$ value |
| --- | --- | --- |
| 4 | 4.4 | <10<sup>-5</sup> |
| 5 | 7.3 | <10<sup>-5</sup> |
| 6 | 12.7 | <10<sup>-5</sup> |
| 7 | 19.0 | <10<sup>-5</sup> |
| 8 | 29.4 | <10<sup>-5</sup> |
| 9 | 28.7 | <10<sup>-3</sup> |
| 10 | 25.3 | <0.01 |
| 11 | 22.7 | <0.01 |
| 12 | 20.1 | <0.05 |

As an alternative means to infer the effect of *ρ*, I recorded the percentage of simulations that retained polymorphism at one or both loci for at least 50 *N_e_* generations across all values of *ρ* (Figure 3). At low values of *N_e_s* (≤ 5), very few polymorphisms (≤ 2.5%) persisted for 50 *N_e_* generations for any value of *ρ*, but there was still a significant effect of *ρ*. At *N_e_s* of 5, a 10-fold reduction in *ρ* more than doubled the number of simulations in which both polymorphisms reached 50 *N_e_* generations (*P* < 10^-5^). At intermediate values of *N_e_s* the effect of *ρ* was quite strong, especially for higher values of *ρ*. At *N_e_s* of 10, fewer than 1% of simulations retained both polymorphisms for 50 *N_e_* generations if *ρ* was greater than or equal to 10, but 17% of simulations did so when *ρ* was 1, an increase of more than 20-fold (*P* < 10^-5^), and a majority of simulations retained both polymorphisms when *ρ* was less than or equal to 0.1. At higher values of *N_e_s*, the effect of *ρ* was again weaker, but still significant. At *N_e_s* of 20, 99% of simulations across all values of *ρ* less than 1 retained polymorphism at both loci for 50 *N_e_* generations, while only 84-94% simulations at values of *ρ* greater than 1 did so (*P* < 10^-3^).

**Figure 3.**
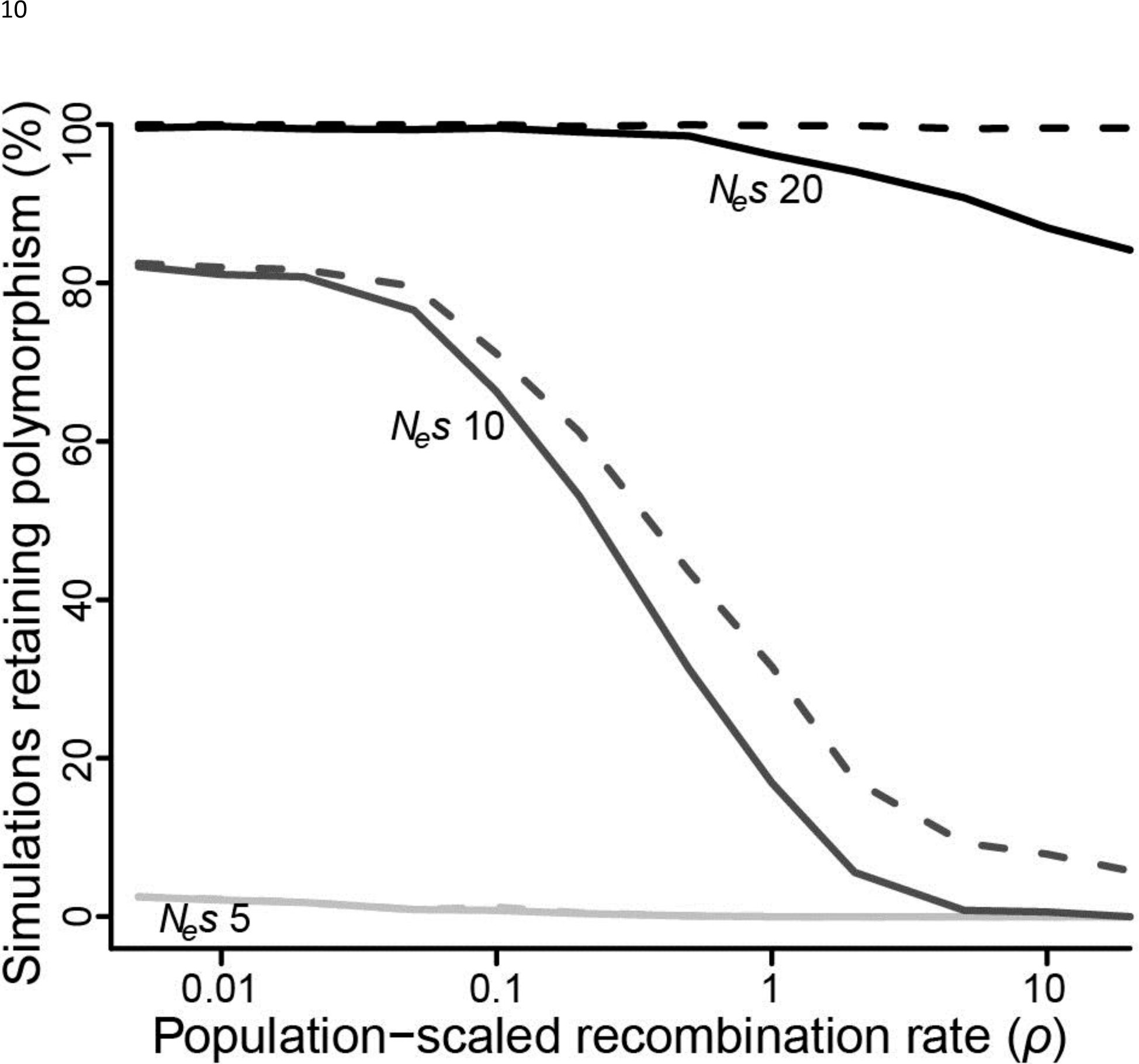
Proportion of simulations where polymorphism is retained after 50 *N_e_* generations, for varying values of selection (*N_e_s* of 5, 10 or 20) and recombination (*ρ* ranging from 0.005 to 20). Solid lines = polymorphism retained at both loci; dashed lines = polymorphism retained at one or both loci.

## Discussion

The fate of a non-neutral allele depends on the strength of selection and the force of genetic drift as determined by the effective population size. When these two evolutionary forces are close to evenly matched, linked polymorphism becomes an important cofactor. A locus under weak balancing selection may or may not retain its polymorphism, depending on whether the purifying effects of genetic drift win out over the effects of selection. However, two or more polymorphisms under weak balancing selection may enjoy a substantial selective advantage if linked to each other, because they can they can mutually boost each other’s adaptive effect. Thus, the combined strength of balancing selection maintaining two or more multi-locus haplotypes could be greater than the effect of either locus on its own. This may especially be true if selection is inconsistent, as polymorphism is likely to be lost during periods when selection does not act on it, unless it is linked to another polymorphism that is being maintained by selection. Similar to a “buddy system” in which two individuals give each other support and protection, pairs or groups of “buddy” polymorphisms can bolster each other. Alleles that would otherwise be swept away by drift are instead anchored by balancing selection. Thus, genetic diversity begets genetic diversity. Polymorphisms maintained for entirely unrelated reasons (e.g. immunity and reproductive strategy) may be more likely to be retained for long periods if they are closely linked.

Putatively balanced polymorphisms are often distributed non-randomly in the genome, but more research is required to empirically document the phenomenon described here. There are several cases where two or more closely-linked loci all show signatures of balancing selection, often with little evidence for epistasis. Some of these are complexes of structurally and functionally related genes, such as immune gene complexes (Takahata and Nei 1990; Takahata et al. 1992; Makino and McLysaght 2008; Tennessen et al. 2015; 2017a) or plant R-gene complexes (Alcázar et al. 2014). In other cases, linked genes appear to have unrelated roles. Population genomic patterns have been most thoroughly characterized in humans, and the relatively low human effective population size (~10^4^, Tenesa et al. 2007) suggests that low values of *N_e_s* and *ρ* may be common in our species. Thus, humans are an obvious system in which to test for linked balanced polymorphisms. For example, human genes *ARPC5* and *RGL1* are less than 1 kb apart, and both show some of the strongest signals of balancing selection in the genome, in populations across multiple continents (DeGiorgio et al. 2014). Unfortunately, it is difficult to discern if such cases are truly independent instances of balancing selection, or the population genetic signature of a single selected polymorphism plus surrounding neutral genetic variation (Siewert and Voight 2017). Balancing selection signatures that are slightly farther apart are less likely to be artifacts of the same balanced polymorphism, but they are also less likely to be mutually supportive. For example, Andrés et al. (2009) identified 60 putative targets of balancing selection across the human genome, including several clusters of seemingly unrelated genes within 300kb of each other (e.g. *LRAP* and *RIOK2; FUT2, PPP1R15A*, and *LHB*). In these cases, distances between linked genes exceed the typical width of linkage disequilibrium blocks in humans (<100kb, Reich et al. 2001), and do not show high linkage disequilibrium in contemporary populations with LDlink (Machiela and Chanock 2015), but are within the range of LD occasionally observed even in panmictic human populations (Koch et al. 2013).

More work is needed to determine the extent to which genomic location influences the fate of an adaptive polymorphism under balancing selection. However, the results presented here suggest that the absence of epistasis is a plausible null hypothesis, and thus tight linkage is not sufficient evidence to conclude that two polymorphic genes interact with each other, even if they share a similar physiological role (e.g. immunological). For example, in *Biomphalaria* snails, the Guadeloupe Resistance Complex consists of several hyperdiverse genes with an apparent role in parasite recognition (Tennessen et al. 2015; Allan and Blouin 2017; Allan et al. 2017a,b), and these are closely linked to another variable locus, *sod1*, also influencing resistance to schistosome parasites (Goodall *et al*. 2006; Tennessen et al. 2017a). While epistasis is an attractive explanation for such clustering of polymorphic, immune-relevant genes, they are not necessarily co-adapted. Similarly, sex chromosomes often undergo dynamic and seemingly adaptive restructuring. Translocation of the sex-determining region has occurred repeatedly in fishes (Lubieniecki et al. 2015), insects (Sharma et al. 2017), and flowering plants (Tennessen et al. 2017b). Such translocations may be favored in order to achieve tight linkage to other polymorphic loci; while such loci are often assumed to be under sexually antagonistic selection and thus in epistasis with the sex locus (Kirkpatrick 2017), such interactions can’t be taken for granted. This may be especially true if dioecy offers little advantage over the hermaphroditic ancestral condition and thus balancing selection acting on the sex locus is relatively weak, as may be the case for wild strawberries (Spigler et al. 2011; Tennessen et al. 2017b).

In summary, many balanced polymorphisms may owe their success to their genomic neighborhood, and not necessarily because of epistasis. Future research characterizing the phenotypic and fitness consequences of tightly linked polymorphisms while be invaluable for documenting and understanding this phenomenon in nature. Are linked adaptive polymorphisms retained by selection more often than solo polymorphisms, and thus disproportionate contributors to segregating adaptive variation? How often are chromosomal rearrangements, such as translocations or inversions, favored by selection because they facilitate tight linkage between functionally unrelated balanced polymorphisms? Is epistasis disproportionately prevalent within genomic regions showing high genetic diversity and/or linkage disequilibrium? The results presented here suggest that linkage *per se* may be an important factor determining distributions and dynamics of adaptive genetic diversity.

## Acknowledgements

I thank M. Blouin, A. Liston, and the Center for Genome Research and Biocomputing At Oregon State University.

